# Single-cell transcriptomics reveals diverse and complex gene expression alterations in human trisomy 18

**DOI:** 10.1101/863332

**Authors:** Jing Wang, Zixi Chen, Fei He, Trevor Lee, Wenjie Cai, Wanhua Chen, Longbin Zhang, Nan Miao, Zhiwei Zeng, Ghulam Hussain, Qiwei Guo, Tao Sun

## Abstract

Trisomy 18, commonly known as Edward’s syndrome, is the second most common autosomal trisomy among live born neonates. Multiple tissues including cardiac, abdominal, and nervous systems are affected by an extra chromosome 18. To delineate the complexity of anomalies of trisomy 18, we analyzed amniotic fluid cells from two normal and three trisomy 18 samples using single-cell transcriptomics. We identified six cell groups, which function in major tissue development such as kidney, vasculature, and smooth muscle, and display significant alterations in gene expression detected by single-cell RNA-sequencing. Moreover, we demonstrated significant gene expression changes in previously proposed trisomy 18 critical regions, and identified three new regions such as 18p11.32, 18q11, 18q21.32, which are likely associated with trisomy 18 phenotypes. Our results indicate complexity of trisomy 18 at the gene expression level and reveal genetic reasoning of diverse phenotypes in trisomy 18 patients.

## INTRODUCTION

Trisomy 18, also known as Edwards syndrome (Edwards et al. 1960), is characterized by the presence of an extra copy of chromosome 18, and is the most common form of autosomal trisomy in live births beside of Down syndrome (trisomy 21) (Carey 2005; Savva et al. 2010; Cavadino and Morris 2017). The incidence of full trisomy 18 is about 94%, compared to all types of trisomy 18 (Cereda and Carey 2012; Mirmohammadsadeghi et al. 2017). Case reports have shown that craniofacial, musculoskeletal, cardiac, abdominal, and nervous systems are most likely to be affected by trisomy 18, even though many organs and tissues display defects (Cereda and Carey 2012; Nagamuthu and Neelaveni 2014; Roberts et al. 2016; Turbiville et al. 2017; Fan et al. 2018). Risk factors for the disease include a positive family history in close relatives and the increasing maternal age (Ginsberg et al. 1968; Calderone et al. 1983; Mirmohammadsadeghi et al. 2017). The prognosis for this genetic defect is poorly understood due to multiple system involvement (Norwitz and Levy 2013; Imataka et al. 2016). Restriction fragment length polymorphisms (RFLP) and short-sequence repeats (SSR) have been exercised to determine the parental origin of the extra chromosome 18 (Kondoh et al. 1988; Kupke and Müller 1989; Nöthen et al. 1993; Ya□gang et al. 1993).

The cause of an extra chromosome 18 is unclear. Studies have shown that nondisjunction predominating at the maternal meiosis I in acroxentric chromosomes results in the addition of an extra chromosome (Eggermann et al. 1996; Bugge et al. 1998). Distal q11.2 at the long arm of chromosome 18 has been proposed as a critical region for the trisomy 18 phenotype, but some controversial views have been reported (Wilson 1993; Boghosian-Sell et al. 1994). The other two critical regions, one within 18q12.1-18q21.2 and one within 18q22.3-18qter, have been proposed (Boghosian-Sell et al. 1994; Roberts et al. 2016). Moreover, a transcriptomic profile of induced pluripotent stem cells from human amniotic fluid cells (AFCs) with trisomy 18 has shown several gene clusters associated with clinical manifestations of trisomy 18 (FitzPatrick et al. 2002; Li et al. 2017; Xing et al. 2018). However, the extremely high mortality rate and complexity of phenotypes in trisomy 18 have hindered genetic and molecular understanding of its etiology (Babu and Verma 1986; Rasmussen et al. 2003; Cereda and Carey 2012; Cavadino and Morris 2017).

The advance of single-cell sequencing technology has allowed comprehensive analyses of gene expression profiles of tens of thousands of single cells at once (Grun and van Oudenaarden 2015; Wen and Tang 2016; Potter 2018). Single-cell sequencing has been used to highlight gene expression in one cell resolution, and to reveal complex and diverse cell types in a tissue and organ (Kim et al. 2018; Park et al. 2018; Raj et al. 2018; Suijuan Zhong1 2018). Thus, it has been expeditiously applied in understanding heterogeneity of organ formation, and complexity of disease conditions (Puram et al. 2017; Young et al. 2018; Zhao et al. 2018; Velmeshev et al. 2019).

In this study, to better understand genetic variations of the trisomy 18 phenotype at one cell resolution, we generate gene expression profiles of amniotic fluid cells from normal and trisomy 18 fetuses using 10X Genomics (Zheng et al. 2017). We show that an extra chromosome 18 has a broad and distinct impact on expression of genes involved in the development of multiple systems such as vascular, renal and nervous systems. We confirm that up-regulation of genes in previously proposed two critical regions (18q12.1-22.1, 18q22.3-18qter) may contribute to the trisomy 18 phenotype. Moreover, we identify three new regions such as 18p11.32, 18q11, 18q21.32, which are likely associated with trisomy 18 defects. Our study demonstrates complexity of trisomy 18 at the gene expression level, and highlights alterations of multiple genes that may contribute to trisomy 18 phenotypes.

## RESULTS

### Single-cell RNA sequencing profiles of trisomy 18 amniotic fluid cells

Trisomy 18 is a hereditary disorder with an extra copy of chromosome 18, which may cause changes in expression levels of many genes, and in turn affects multiple organs (Antonarakis et al. 2004; Antonarakis 2017). To understand complexity of such anomalies, we analyzed single-cell transcriptomics of amniotic fluid cells associated with normal and trisomy 18 fetuses using 10X Genomics. This approach allowed bypass of direct fetal tissue collection and facilitated deciphering diversity in amniotic fluid cells from trisomy 18 fetuses, which may reflect complexity of this disorder.

Human amniotic fluid cells were collected from amniocentesis screen for hereditary disorders using the karyotype analysis. Based on karyotypes, we selected amniotic fluid samples from two healthy and three trisomy 18 fetuses for single cell transcriptomics analyses (Supplemental Fig. S1A-C). We sequenced and analyzed 56,517 cells, in which 18,281 cells were from normal samples and 38,236 cells from trisomy 18 (Supplemental Table S1). We further grouped these cells into 13 transcriptionally distinct clusters using non-linear dimensionality reduction with spectral t-distributed stochastic neighbor embedding (tSNE) (Supplemental Fig. S1D). In details, two normal samples were grouped into 11 clusters, while three trisomy 18 samples were grouped into 10, 8 and 10 clusters, respectively (Supplemental Fig. S1E-H). These results indicate that amniotic fluid cells from normal and trisomy 18 fetuses display a diverse transcriptional identity.

To further classify major cellular functions, we annotated genes from the 13 clusters according to their major functions using gene ontology (GO) analyses (Supplementary Fig. S2A, Supplementary Table S2). We identified 6 functional categories, and subsequently classified two normal samples into 6 sub-clusters, which play a role in kidney function, vasculature development, smooth muscle cell proliferation, protein synthesis, nuclear division and metabolic process (Fig. 1A, B). Trisomy 18 samples were grouped into similar 6 functional sub-clusters, with different cell numbers in each sub-cluster. For example, compared to normal individuals, trisomy 18 sample-1 (T18-1) displayed very few cells with the function in nuclear division and protein synthesis, T18-2 and T18-3 had fewer cells in metabolic process (Fig. 1B, C and Supplementary Fig. S2C, D). These data suggests that trisomy 18 amniotic fluid cells display similar functional groups but altered cell components in each group, compared to those in normal ones.

**Figure 1.**
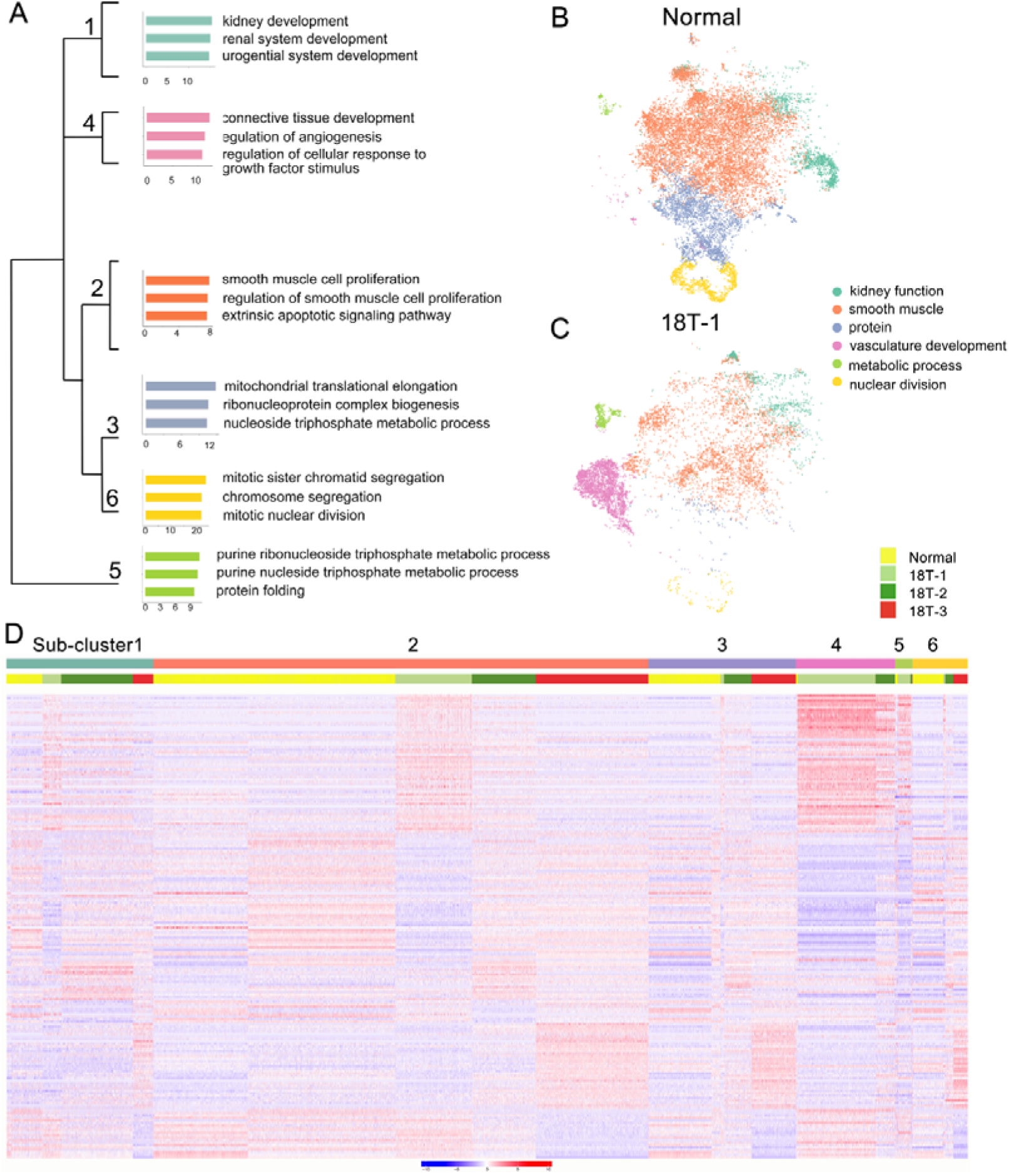
Functional grouping of amniotic fluid cells from normal and trisomy 18 (T18) samples. (*A*) Different functional sub-clusters were annotated according to expression of significantly altered genes for two normal samples. Gene ontology (GO) analyses of differentially expressed genes in three trisomy 18 samples were compared with two normal samples. (*B, C*) t-distributed Stochastic Neighbor Embedding (t-SNE) projection of cells and typical gene expression in normal and T18-1 cells. (*D*) Heatmap of differentially expressed genes in two normal and three trisomy 18 samples.

To reveal gene expression patterns in 6 sub-clusters, we randomly selected 2,000 cells per sub-clusters, and top 100 differentially expressed genes in each sub-cluster by using a random forest classification. Compared to normal samples, genes in each trisomy 18 sample displayed either up- or down-regulated expression in each functional sub-clusters (Fig. 1D). For example, the sub-cluster 4 was functionally grouped as vasculature development, and genes in T18-1 and T18-2 displayed higher expression in this sub-cluster. Moreover, in each sub-cluster, genes in T18-3 were also noticed with a differential expression compared to those in T18-1 and T18-2. For instance, compared to normal samples, genes grouped in the sub-cluster 1 showed up-regulation in T18-1, -2 and -3, but the up-regulated genes were distinct in the T18-3 sample. These results suggest that an extra copy of chromosome 18 causes altered gene expression in human amniotic fluid cells, and this alteration is distinct in each sample.

### Differentially expressed genes in trisomy 18 amniotic fluid cells

Studies have shown that multiple organs are affected in trisomy 18 (Cereda and Carey 2012). We next wanted to identify major genes and pathways that might be altered in each functional sub-cluster at a single cell level. According to numbers of cells, detected genes (P<0.01), and unique molecular identifiers (UMIs) in every sub-clusters, we identified 2,523 specific genes in 6 sub-clusters (Fig. 2A, and Supplementary Table. S3). Representative genes in each sub-clusters were identified, for example *STAT1, DLG1,* and *EDN1* in the sub-cluster-1 of kidney development, *IGFBP2* and *PDGFB* in the sub-cluster-2 of smooth muscle cell proliferation, *VEGFA, VEGFC, FLT1,* and *ZNF703* in the sub-cluster-4 of regulation of vascularization, and *CDC20, PRC1, NUSAP1,* and *CCNB1* in the sub-cluster-6 of mitotic nuclear division (Fig. 2B).

**Figure 2.**
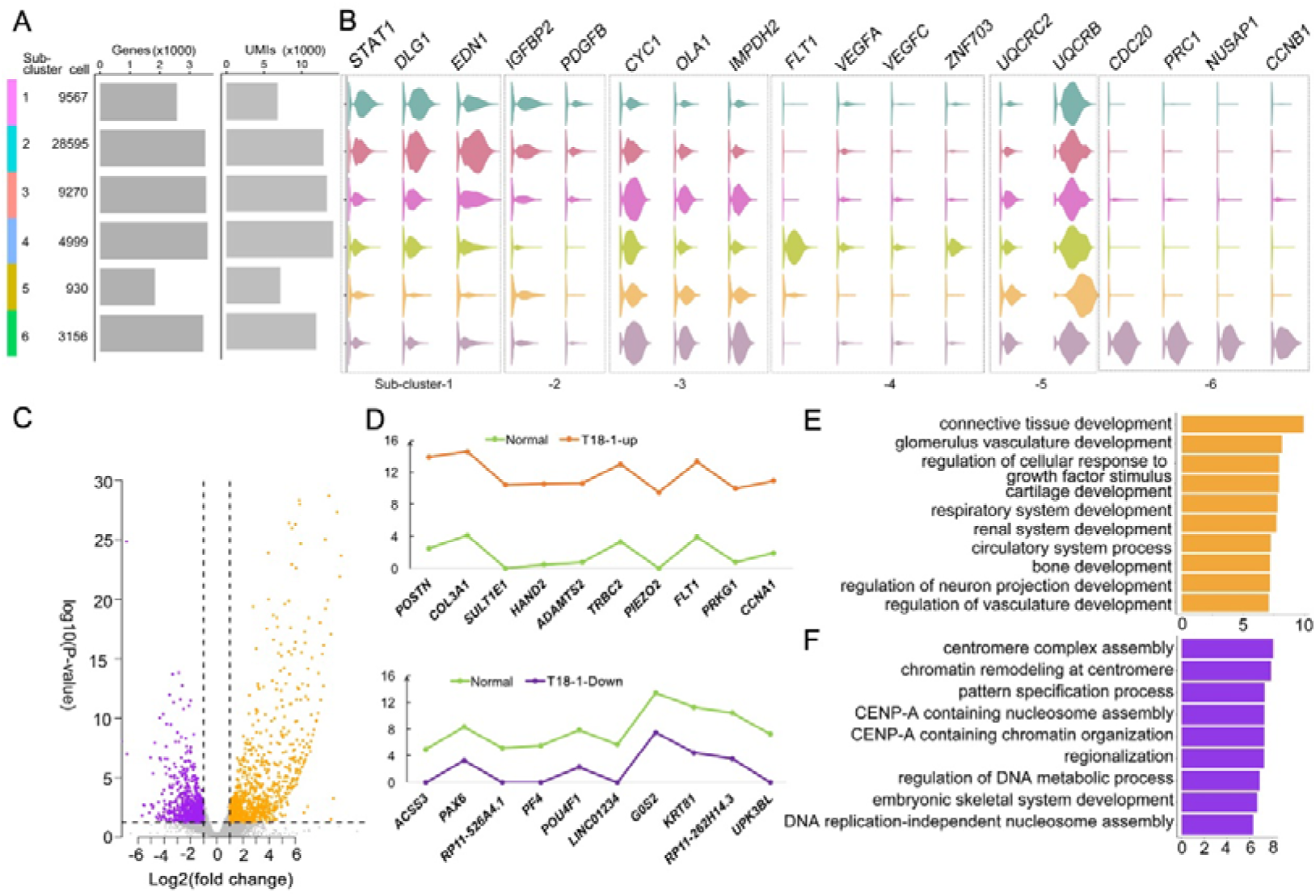
Altered genes in trisomy 18 amniotic fluid cells. (*A*) Dendrogram of 6 sub-clusters, cell number per sub-cluster, number of genes in each sub-cluster (P<0.01), and unique molecular identifiers (UMIs). (*B*) Violin plots of representative genes identified in 6 sub-clusters. (*C*) Volcano plot of differentially expressed genes by comparing the T18-1 with normal samples. Up-regulated genes: orange dots; down-regulated genes: purple dots. (*D*) Top 10 (-log2fold change>1 and FDR corrected P≤0.05) up-regulated genes (orange), down-regulated genes (purple) in the T18-1 sample compared with two normal samples (green). (*E*) Gene ontology (GO) analysis of up-regulated genes in T18-1 cells compared with two normal samples. (*F*) GO analysis of down-regulated genes in 18T-1 cells compared with two normal samples.

We then compared differentially expressed genes (P≤0.01) between every trisomy 18 sample and two normal samples (Fig. 2C, and Supplementary Fig. 3). In the T18-1 sample, a majority of differentially expressed genes was up-regulated (Fig. 2C). Among all differentially expressed genes (-log2fold change>1 and FDR corrected P≤0.05), top 10 up-regulated and down-regulated genes in T18-1 cells were listed (Fig. 2D and Supplementary Table S4). Moreover, we performed GO analyses for differentially expressed genes in T18-1 cells. GO terms for up-regulated genes were enriched in regulation of glomerulus vasculature development, circulatory system development, and neuron projection development (Fig. 2E). GO terms for down-regulated genes were enriched in regulation of centromere complex assembly, pattern specification, and DNA metabolic process (Fig. 2F and Supplementary Table S5, 6). These analyses suggest that an extra chromosome 18 alters expression of genes that are associated with major organ developmental processes and fundamental cellular functions.

Furthermore, we analyzed and compared differentially expressed genes in T18-2 and T18-3 samples with two normal samples (Supplementary Fig. 3A, B and Supplementary Table S7, 8). Relative expression levels of top 10 up-regulated and down-regulated genes from T18-2 and T18-3 samples were listed (Supplementary Fig. 3C, D). Interestingly, *POSTN*, *COL3A1*, *FLT1*, *ADAMTS2* were up-regulated in both T18-1 and T18-2 cells, while *UPK3BL* was down-regulated in both T18-1 and T18-3 cells (Fig. 2 and Supplementary Fig. 3C, D).

Finally, the GO term of renal system development was enriched in up-regulated genes from all three trisomy 18 samples (Supplementary Table S9, 11), while neuron fate commitment was enriched for down-regulated genes in 18T-2 and 18T-3 (Supplementary Table S10, 12). In addition, among up-regulated genes, the GO term of stem cell development was enriched in T18-3 cells. And among down-regulated genes, GO terms of neuron fate commitment, negative regulation of oligodendrocyte differentiation, peripheral nervous system neuron development and spinal cord motor neuron cell fate specification were enriched in the T18-2 sample, and they were enriched in kidney development and embryonic digestive tract development in the T18-3 sample (Supplementary Fig. 3E, F). These results further indicate that while the extra chromosome 18 has a broad impact on development of renal and nervous systems, it also preferentially affects different genes associated with either stem cells, neurons, or digestive track cells in each trisomy 18 sample.

### Trisomy 18 alters expression of genes located on all chromosomes

We further investigated the effect of an extra copy of chromosome 18 on expression of genes located on all chromosomes. We merged expression levels of all genes detected by single-cell sequencing on each chromosome, and compared their differential expression in normal and trisomy 18 samples using Deseq2. Box plots showed increased expression of genes located on chromosome 18 in trisomy 18 samples (Fig. 3A). Moreover, gene expression levels (-log10P-value) in each chromosome also showed an up-regulated gene expression in trisomy 18 samples (Fig. 3B). These data indicates that an extra chromosome 18 indeed causes up-regulated expression of genes located on chromosome 18.

**Figure 3.**
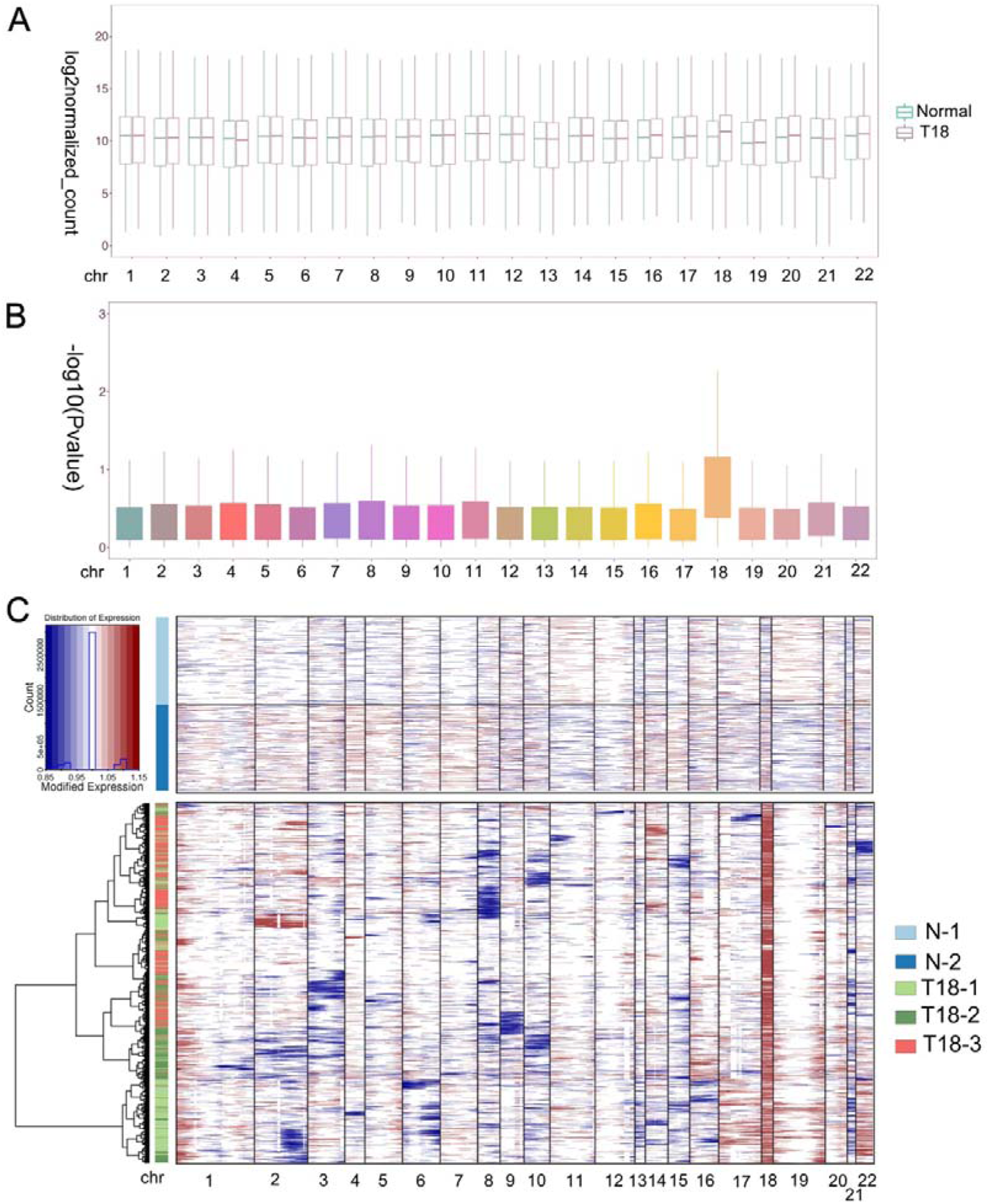
Trisomy 18 alters gene expression on chromosomes. (*A*) Merged expression levels of genes, showing log2 normalized counts, on each chromosome. (*B*) Up-regulated gene expression (-log10p-value) on chromosome 18 from trisomy 18 samples. (*C*) Heatmap of large-scale chromosomal copy-number variations (CNVs). Normal cells served as reference cells, and trisomy 18 samples (T18-1, T18-2 and T 18-3) as observed cells.

We next analyzed single-cell transcriptomics data including gene numbers and expression levels, based on randomly selected 2,000 cells in each normal and trisomy 18 samples. We inferred large-scale chromosomal copy-number variations (CNVs) in every single cell that is based on averaged expression profiles across chromosome intervals (Patel et al. 2014; Muller et al. 2016; Filbin et al. 2018). CNV analyses and inference of haplotypes uncovered distinct genetic variations in trisomy 18 samples, and CNVs in each trisomy 18 samples showed distinct alterations (Fig. 3C, Supplementary Fig. 4). As a consequence of the third chromosome 18, CNVs were greatly increased in trisomy 18 samples (Fig. 3C). In addition, CNVs from chromosome 1, 19 and 20 showed an increase, and those from chromosome 3, 8, 9, 10, 15 and 21 displayed a decrease in trisomy 18 samples (Fig. 3C and Supplementary Fig. 4B-D). These results suggest that the third chromosome 18 affects the expression of genes on all chromosomes, and causes altered copy numbers of genes on them.

### Gene expression patterns in proposed trisomy 18 critical regions

Previous studies proposed a few chromosomal regions as critical regions for trisomy 18 phenotypes, such as 18q12.1-21.2 and 18q22.3-18qter (Boghosian-Sell et al. 1994). We thus examined whether altered expression of genes within these regions might be responsible for trisomy 18 phenotypes.

The human chromosome 18 consists of 984 genes (Homo sapiens. GRCh38). Our single-cell transcriptomics identified 292 up-regulated genes out of 984 genes in all trisomy 18 samples (P< 0.05) (Supplementary Table. S13). Within the proposed 18q12.1-21.2 critical region, single-cell transcriptomics identified 78 genes out of 292 genes (Fig. 4A). Among these 78 genes, 58 genes were significantly up-regulated (P<0.05) and displayed higher copy numbers detected by the CNV analysis (Fig. 4B, C). The up-regulated genes were enriched in ventricular cardiac muscle tissue development and mesonephric tubules in the GO term analysis (Fig. 4G).

**Figure 4.**
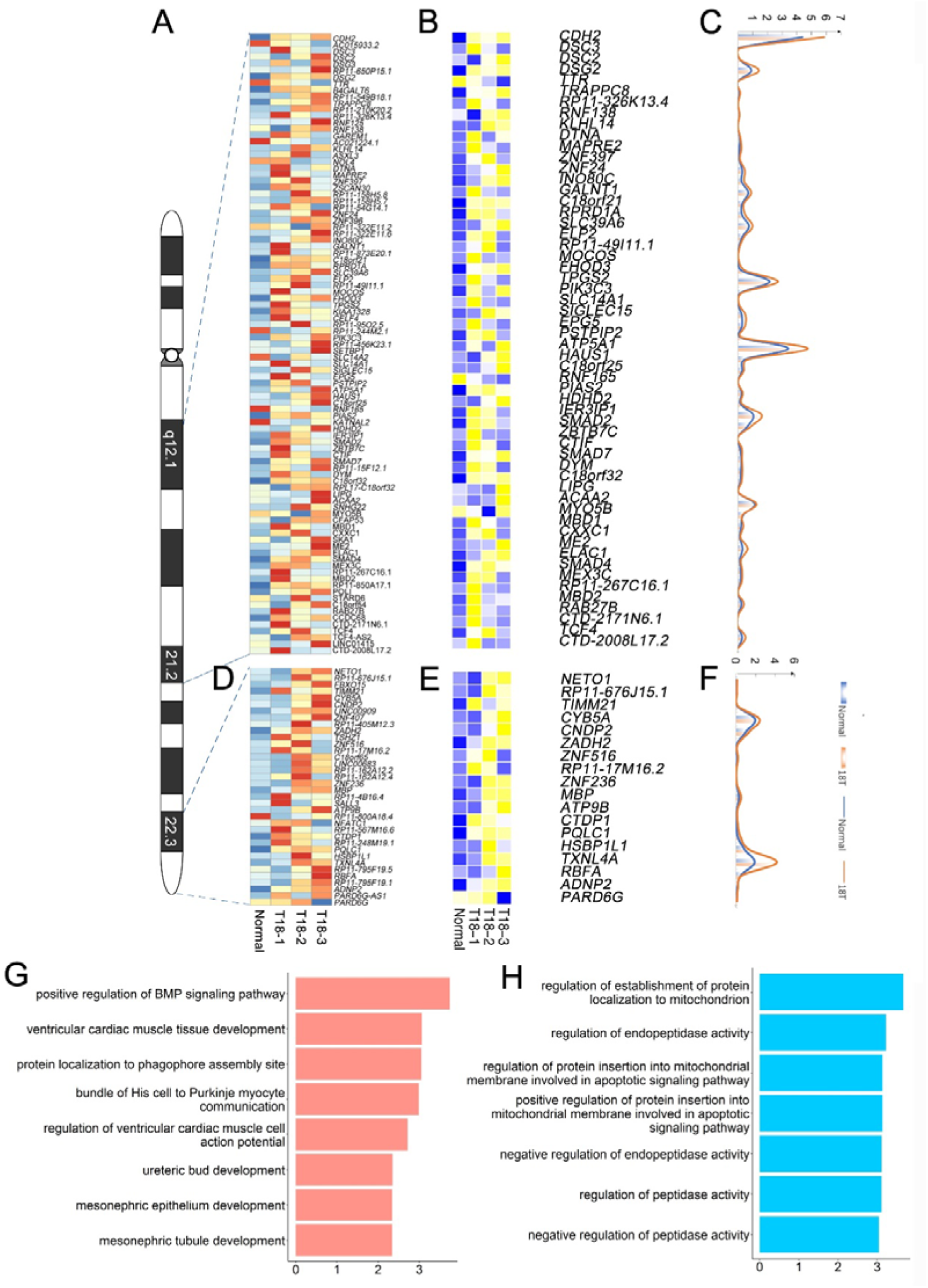
Gene expression analyses in previously proposed trisomy 18 critical regions: 18q12.1-21.2 and 18q22.3-18qter. (*A*) Heatmap of expression levels of genes in the 18q12.1-21.2 region from two normal and three trisomy 18 samples. (*B*) Heatmap of differentially expressed genes (*p* ≤0.01) in the 18q12.1-21.2 region. (*C*) Unique molecular identifiers (UMIs) counts of differentially expressed genes in normal (blue line) and trisomy 18 samples (orange line). (*D*) Heatmap of expression levels of genes in the 18q22.3-18qter region from two normal and three trisomy 18 samples. (*E*) Heatmap of differentially expressed genes (*p* ≤0.01) in the 18q22.3-18qter. (*F*) UMIs counts of differentially expressed genes in normal (blue line) and trisomy 18 samples (orange line). (*G*) Top gene ontology (GO) terms of differentially expressed genes in the 18q12.1-21.2 region. (*H*) Top GO terms enriched for differentially expressed genes in the 18q22.1-18qter region.

Moreover, 110 genes were identified in the proposed 18q22.3-qter critical region. Among these 110 genes, single-cell transcriptomics identified 36 genes with altered expression (Fig. 4D). Of them, 18 genes showed a significantly up-regulated expression (P <0.05), and a higher copy number detected by the CNV analysis (Fig. 4E, F). Up-regulated genes were enriched in regulation of establishment of protein localization to mitochondrion and negative regulation of endopeptidase activity in the GO term analysis (Fig. 4H).

Overall, our results indicate that single-cell transcriptomics confirms the previously reported critical regions, and identifies crucial genes that are involved in development of cardiac muscle and mesonephric tubule, and in basic cellular function such as protein localization.

### Prediction of new chromosome 18 critical regions

Because of the complexity of phenotypes in trisomy 18 patients, we suspected that there might be some other trisomy 18 critical regions, which contribute to these defects. Based on the sequencing data of single-cell transcriptomics, we screened out differentially expressed genes (P<0.05) in all five normal and trisomy 18 samples by DESeq2. We merged all differentially expressed genes from trisomy 18 samples, and mapped them onto chromosome 18. We identified three regions, in which genes are significantly up-regulated (P<0.05) and have higher copy numbers detected by the CNV analysis. These regions included 18p11.32 on the short arm, and 18q11 and 18q21.32 on the long arm of chromosome 18.

Single-cell transcriptomics showed that 15 genes out of 57 genes on 18p11.32, 33 genes out of 101 genes on 18q11, and 12 genes out of 47 genes on 18q21.32 were significantly up-regulated (Fig. 5A-C). The degree of up-regulation varied among these genes. Moreover, in the 18p11.32 region, up-regulated genes were enriched in negative regulation of mitotic metaphase plate congression, centrosome duplication, and nuclear division in the GO term analysis (Fig. 5D). In the 18q11 region, up-regulated genes were enriched in endodermal cell differentiation, in utero embryonic development, and cardiac muscle hypertrophy (Fig. 5E). Finally, in the 18q21.32 region, up-regulated genes were enriched in protein localization to mitochondrion, lymphangiogensis, and positive regulation of T cell cytokine production (Fig. 5F). These analyses suggest that these new regions in chromosome 18 might contribute to the complexity of phenotypes in trisomy 18 patients, including cardiac muscle development, immune response and basic cell division regulation.

**Figure 5.**
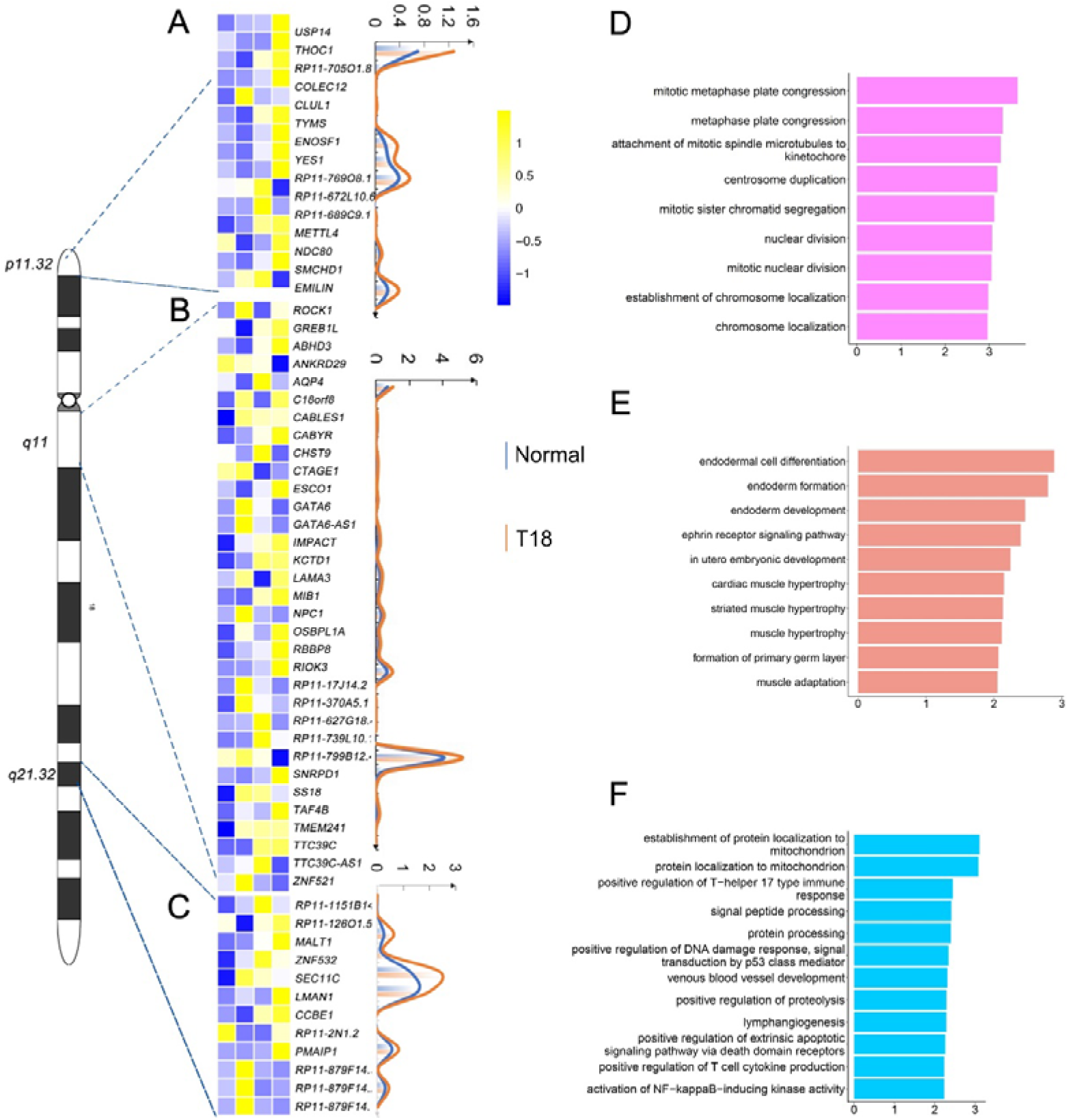
Gene expression analyses in 18p11.32, 18q11 and 18q11.32 regions. (*A*) Heatmap of differentially expressed genes (*p* ≤0.01) in the 18p11.32 region from two normal and three trisomy 18 samples. Unique molecular identifiers (UMIs) counts of differentially expressed genes in normal (blue line) and trisomy 18 samples (orange line). (*B*) Heatmap of differentially expressed genes (*p* ≤0.01) in the 18q11 region. UMIs counts of differentially expressed genes in normal (blue line) and trisomy 18 samples (orange line). (*C*) Heatmap of differentially expressed genes (*p* ≤0.01) in the18q21.32 region. UMIs counts of differentially expressed genes in normal (blue line) and trisomy 18 samples (orange line). (*D-F*) Top gene ontology (GO) terms of differentially expressed genes in the 18q11.32 (*D*), 18p11 (*E*) and 18q21.32 (*F*) regions.

## DISCUSSION

Trisomy 18 pregnancies, characterized by an extra copy of chromosome 18, have a high risk of fetal loss and stillbirth (Won et al. 2005; Morris and Savva 2008). Insightful studies on pathophysiology and genetic causes of trisomy 18 have been sparse, because of the difficulty of sample collection, unavailability of disease models and extremely high mortality rate among neonates (Cavadino and Morris 2017; Xing et al. 2018). Here, we have analyzed three amniotic fluid cell samples from trisomy 18 fetuses using the single-cell sequencing technology. We find that the extra chromosome 18 alters expression of genes functioning in multiple systems such as vascular, renal and nervous systems. We demonstrate striking gene expression changes in previously proposed trisomy 18 critical regions, and identify three new regions such as 18p11.32, 18q11, 18q21.32, which are likely associated with trisomy 18 phenotypes. Our study indicates complexity of trisomy 18 at the gene expression level revealed by the single-cell sequencing.

Trisomy 18 is detected antenatally in 43% cases, and almost 90% of those born without a diagnosis is known to be growth retarded *in utero* (Embleton et al. 1996). A decrease in the frequency of live born with trisomy 18 has been observed because of increased use of prenatal diagnosis and the high rate of pregnancy termination based on such diagnosis (Crider et al. 2008; Irving et al. 2011). The amniotic fluid has been collected as an untapped source of cells with broad potential, such as prenatal diagnosis and genetic analysis, which don’t have the ethical and legal limitations of embryonic stem cells (De Coppi et al. 2007; Loukogeorgakis and De Coppi 2016; Loukogeorgakis and De Coppi 2017). In this study, we thus use amniotic fluid cells as a source to study complexity of the trisomy 18 phenotype based on gene expression. While RNA-sequencing has been widely used to generate gene expression profiles in a tissue and organ, it cannot distinguish diversity and complexity of cell composition. Single-cell sequencing can reveal comprehensive transcriptomics at an individual cell resolution, overcome limited tissue supply, identify novel cell types, and uncover diverse contributions of individual cells to organ formation and disease conditions (Tang et al. 2009; Tang et al. 2010; Haque et al. 2017; Hochgerner et al. 2018; Li et al. 2018; Potter 2018). Therefore, in this study, we have applied the single-cell sequencing approach to examine the composition of amniotic fluid cells from normal and trisomy 18 fetuses.

Clinical studies indicate that among trisomy 18 phenotypes, 75% cases show malformations in heart, 25%-75% in genitourinary system, and 5-25% in gastrointestinal system, central nervous system, craniofacial, eye and limb, suggesting that multiple systems are affected by the extra chromosome 18 (Hodes et al. 1978; Lyons Jones 1988; Balderston et al. 1990; Root and Carey 1994; Cereda and Carey 2012; Hoon et al. 2018). Our single-cell sequencing has identified 6 functional sub-clusters, in particular cell groups functioning in the development of kidney, vasculature, and smooth muscle, which display significant alterations in gene expression. Our finding has provided genetic reasoning that is associated with multiple and as well specific trisomy 18 defects. Moreover, we have found that up- and down-regulated genes are similar but also distinct among three trisomy 18 samples. While variations of amniotic fluid cells from three individual fetuses may contribute to diversity of altered genes, distinct alterations of gene expression in each single cells in three individuals, which has been captured by single-cell sequencing, is likely associated with the complexity of trisomy 18 defects. Nevertheless, our GO term analyses indicate that up-regulated genes detected in three trisomy 18 samples are commonly enriched in development of the renal system, bone, and urogenital system. Single-cell sequencing of amniotic fluid cells from trisomy 18 fetuses not only reveals diverse gene expression programs in each cell from each individual, but also identifies common genes that are associated with major defects in trisomy 18.

Identifying critical regions in a chromosome is an effective way to uncover genetic reasoning of autosomal trisomy disorders (Mrozek et al. 2002; Olson et al. 2004; Chang and Min 2005; Li et al. 2014; Mowery et al. 2018). 18q12.1-18q21.2 and 18q22.33-18qter have been proposed as trisomy 18 critical regions (Boghosian-Sell et al. 1994). We have found that genes in these two regions are indeed significantly up-regulated in trisomy 18 amniotic fluid cells. Moreover, GO analyses of up-regulated genes also support functional association with cardiac ventricular development and protein localization, which highlights tight relation of these two regions with trisomy 18 defects. Furthermore, because of the involvement of multiple tissues in trisomy 18 anomalies, it appears unlikely only two trisomy 18 critical regions contribute to the phenotypes. Our results suggest that three new regions (18p11.32, 18q11 and 18q21.32) are tightly associated with trisomy 18 defects. Expression levels of genes located in these regions are significantly altered, and these genes are functionally involved in cell division regulation, endoderm formation, and immune response. Our finding indicates that trisomy 18 affects immune system and fundamental cellular function such as cell division, and in turn has a broad impact on proper development and function of multiple systems.

Overall, our single-cell sequencing analysis reveals complexity of gene expression alterations in amniotic fluid cells from trisomy 18 fetuses. The previously proposed and newly identified trisomy 18 critical regions highlight genetic contributions to the aggravation of symptoms in the trisomy 18 condition.

## METHODS

### Amniotic fluid cells acquisition, culturing and karyotype analyses

Amniotic fluid cell samples were recruited from pregnant women who underwent amniocentesis for prenatal diagnoses at Women and Children’s Hospital, School of Medicine, Xiamen University. Signed informed consents had been obtained from pregnant women who received amniocentesis for prenatal diagnosis to agree the research use of the remainder amniotic fluid samples. The culture cells used in this study were those remainder samples. There was no additional sampling or any health-related intervention performed. Moreover, except for karyotypes, other information including names of patients was withheld from the study group.

Cells were cultured according to standard protocols of using the flask method and commercially available complete medium for amniotic fluid cells (Amniopan, Biotech) at 37°C in 5% CO_2_ environment. The karyotype analysis was conducted according to standard protocols for the chromosomal Giemsa (G)-banding. A total of 100 metaphases of each passage was evaluated.

Following a routine cytogenetic analysis, the second passage of amniotic cell culture was grown in the same condition as for the primary cell culture, and was used for single-cell sequencing.

### Preparation of single-cell suspension

Cultured amniotic fluid cells were digested with 0.05% trypsin-EDTA (Gibco, Life Technologies) at 37°C for 3 minutes. The reaction was terminated by adding the complete medium. Cell suspension was pipetted to dissociate it into single cells, and centrifuged at 1000 rpm for 5 minutes to obtain a cell pellet. The cell pellet was suspended in 5 ml of Phosphate Buffered Saline (PBS), and was centrifuged again at 1000 rpm to remove cell debris. The cell pellet was re-suspended into single cells in 100 μl ice-cold PBS for single-cell sequencing analysis.

### Single-cell RNA sequencing (ScRNA-seq) library construction using the 10X genomics chromium platform

ScRNA-seq libraries were prepared per the Single Cell 30 Reagent Kit User Guide v2 (10X Genomics). Briefly, cell suspensions were loaded on a Chromium Controller instrument (10X Genomics) to generate single-cell Gel Bead-In-EMulsions (GEMs). GEM-reverse transcriptions (GEM-RTs) were performed in a Veriti 96-well thermal cycler (Thermo Fisher Scientific). After reverse transcription, GEMs were harvested, and cDNAs were amplified and cleaned up with the SPRIselect Reagent Kit (Beckman Coulter). Indexed sequencing libraries were constructed using the Chromium Single-Cell 30 Library Kit (10X Genomics) for enzymatic fragmentation, end-repair, A-tailing, adaptor ligation, ligation cleanup, sample index PCR, and PCR cleanup. The barcoded sequencing libraries were quantified by quantitative PCR using the KAPA Library Quantification Kit (KAPA Biosystems). Sequencing libraries were loaded on a NextSeq500 (Illumina) with a custom sequencing setting (26 bp for read 1 and 98 bp for read 2) to obtain a sequencing depth of 80,000 reads per cell (Xie et al. 2018).

### Data processing of single-cell RNA-seq from the chromium system

Mapping to the GRCh38 human genome, quality control and read counting of Ensembl genes were performed using Cellranger software with default parameter (v2.1.0). Cells with ≥200 genes and ≤ 10% percentage of detected mitochondrial genes were retained for subsequent analyses. Normalization, dimensionality reduction and clustering of single cells were also performed by the Cellranger.

### Identification of differentially expressed genes

Differentially expressed genes (DEGs) were first identified by comparing cells in a specific cluster with cells in all other clusters (Seurat package Wilcox: average expression difference > 0.5 natural log with an FDR corrected P<0.01). Next, cells in a specific cluster were compared to cells in every other cluster in a pairwise manner to identify a second set of DEGs (DESeq2 package: FDR ≤ 0.05 and |log2 Fold Change| >= 2).

### Identification of cell types and subtypes by nonlinear dimensional reduction and random forests

Batch effects were corrected by matching mutual nearest neighbors (Haghverdi et al. 2018), and the dimensionality was reduced by running a principal-component analysis (PCA) on the most highly variable genes. Graph-based clustering was performed on significant principal components (PCs).

The Seurat package (v.3.0.0) implemented in R was applied to identify major cellular functions of 56,517 single cells from amniotic fluid cells. The top 1,500 most variable genes selected by Seurat were used to compute the PCs. The first 50 significant (p≤0.01) PCs selected based on the built-in jackstraw analysis were used for t-distributed stochastic neighbor Embedding (t-SNE) visualization. Dimensionality was reduced by running a PCA on the most highly variable genes, and the graph-based clustering was performed on the significant PCs. Finally, the distinct sub-groups of cells were visualized by using t-distributed stochastic neighbor embedding tSNE.

### Analysis of copy-number variations (CNVs) in single-cell data

Analysis of CNVs with single-cell data was conducted using the R package inferCNV (https://github.com/broadinstitute/inferCNV) with default parameters.

## DATA ACCESS

Public database accession numbers: GSE141074 provides access to all data. (https://www.ncbi.nlm.nih.gov/geo/query/acc.cgi?acc=GSE141074)

## ACKNOELEDGMENTS

This work was supported by the Scientific Research Funds of Huaqiao University (Z17Y0026 and Z16Y0017), Innovation Awards of Quanzhou Talents (2018C057R), Natural Science Foundation of Fujian Province, China (2019J01071), China Postdoctoral Science Foundation (2017M622053), and National Natural Science Foundation of China (31771141).

